# Formylation facilitates the reduction of oxidized initiator methionines

**DOI:** 10.1101/2024.02.06.579201

**Authors:** Ruiyue Tan, Margaret Hoare, Philip Bellomio, Sarah Broas, Konttessa Camacho, Kyle Swovick, Kevin A. Welle, Jennifer R. Hryhorenko, Sina Ghaemmaghami

## Abstract

Within a cell, protein-bound methionines can be oxidized by reactive oxygen species (ROS) or monooxygenases, and subsequently reduced by methionine sulfoxide reductases (Msrs). Methionine oxidation can result in structural damage or be the basis of functional regulation of enzymes. In addition to participating in redox reactions, methionines play an important role as the initiator residue of translated proteins where they are commonly modified at their α-amine group by formylation or acetylation. Here, we investigated how formylation and acetylation of initiator methionines impact their propensity for oxidation and reduction. We show that *in vitro*, N-terminal methionine residues are particularly prone to chemical oxidation, and that their modification by formylation or acetylation greatly enhances their subsequent enzymatic reduction by MsrA and MsrB. Concordantly, *in vivo* ablation of methionyl-tRNA formyltransferase (MTF) in *E. coli* increases the prevalence of oxidized methionines within synthesized proteins. We show that oxidation of formylated initiator methionines is detrimental in part because it obstructs their ensuing deformylation by peptide deformylase (PDF) and hydrolysis by methionyl aminopeptidase (MAP). Thus, by facilitating their reduction, formylation mitigates the misprocessing of oxidized initiator methionines.

Classification: Biological Sciences; Biochemistry

## Introduction

Methionine (Met) residues are highly prone to oxidation and are readily converted to methionine sulfoxides (MetO) through the reaction of their sulfur-containing sidechains with reactive oxygen species (ROS)^1–3^. Because oxidation alters the hydrophobic nature of methionine residues, the conversion of Met to MetO is often structurally deleterious and results in protein misfolding and aggregation^4^. Accordingly, methionine oxidation has been associated with a number of proteins misfolding disorders and pathological aging^1,5,6^. MetO residues can be enzymatically reverted to Met through the action of a conserved class of enzymes known as methionine sulfoxide reductases (Msrs)^7,8^. Distinct families of Msrs stereo-selectively reduce MetO in either protein-bound or free forms. For example, the three well-studied Msrs in *E. coli* - MsrA, MsrB and MsrC (fRMsr) - are primarily responsible for reduction of protein-bound methionine-S-sulfoxides, protein-bound methionine-R-sulfoxides, and free methionine-R-sulfoxides, respectively^9,10^.

Historically, formation of MetO has been classified as a form of oxidative damage and Msrs were primarily thought of as protein repair enzymes. However, recent studies have shown that reversible methionine oxidation is the regulatory basis of a number of enzymes^11–14^. Additionally, it has been shown that continuous cycles of chemical oxidation and enzymatic reduction of methionines can act as a ROS scavenger mechanism within cells, preventing the oxidation of other biomolecules and mitigating the effects of oxidative stress^15–17^. Thus, the functional ramifications of methionine oxidation extend beyond stochastic structural damage.

In addition to their important redox properties, methionines also play a critical role in protein synthesis. Within both prokaryotic and eukaryotic cells, protein translation is initiated with methionine residues^18^. In prokaryotes, as well as mitochondria and chloroplasts of eukaryotes, initiator methionines are modified by the addition of a formyl group to their α-amine group^19^. The formyl group is enzymatically added to methionyl-tRNAs by methionyl-tRNA formyltransferase (MTF) prior to formation of the translation initiation complex^20^. The formyl modification is subsequently removed by peptide deformylase (PDF) during translational elongation^21^. In the eukaryotic cytosol, the α-amine group of some initiator methioinines can be analogously modified by addition of an acetyl group by *N*-terminal acetyltransferases (NATs)^22^. In both prokaryotes and eukaryotes, a subset of initiator methionines are retained by mature proteins, whereas others are removed from nascent polypeptides through the action of methionine aminopeptidases (MAPs)^23^.

The functional significance of α-modification of methionines is incompletely understood^24^. It has been shown that α-modifications of methionines can impact the synthesis, folding, binding properties and degradation of proteins^25,26^. In bacteria, although methionine formylation is known to facilitate translational initiation, it is not strictly essential for either protein synthesis or survival^20,24,27^. Nonetheless, recent studies suggesting that formylated proteins are selectively targeted for degradation by the N-degron pathways provide important clues regarding the possible functional significance of formylation^24,28^. Although methionine formylation is typically considered to be a prokaryotic modification, it is becoming increasingly evident that it also plays an important role in eukaryotic biology. In humans, defects in the formylation and deformylation of mitochondrial proteins have been linked to apoptosis, mitochondrial disfunction and disorders such as Leigh syndrome^29–31^. Furthermore, it has been shown that methionine formylation can occur in eukaryotic cytosols as a response to cellular stress^31^. Despite these recent findings, the functional logic of transiently formylating initiator methionines within nascent polypeptides remains enigmatic.

Although the oxidation and formylation of methionine residues have been individually well studied, it is not known how these two important modifications influence each other. Here, we have sought to investigate how formylation impacts the redox properties of initiator methionines, and conversely, how the oxidation status of initiator methionines affects their formylation and subsequent processing in nascent polypeptides.

## Results

### Formylation facilitates reduction of Nt-methionine by Msrs in a synthetic peptide

To determine whether formylation impacts the redox kinetics of methionines, we initially conducted *in vitro* oxidation and reduction experiments on methionine-containing synthetic peptides (Figure 1). The analyzed peptides contained N-term methionine residues that were either unmodified (Nt-Met) or formylated (Nt-fMet). The peptides were oxidized with varying concentrations of hydrogen peroxide and fractional oxidation levels of methionines were subsequently quantified by mass spectrometry (Figure S1). The results indicated that Nt-fMet has a faster oxidation rate than Nt-Met, although the magnitude of this difference was slight (Figure 1A).

**Figure 1.**
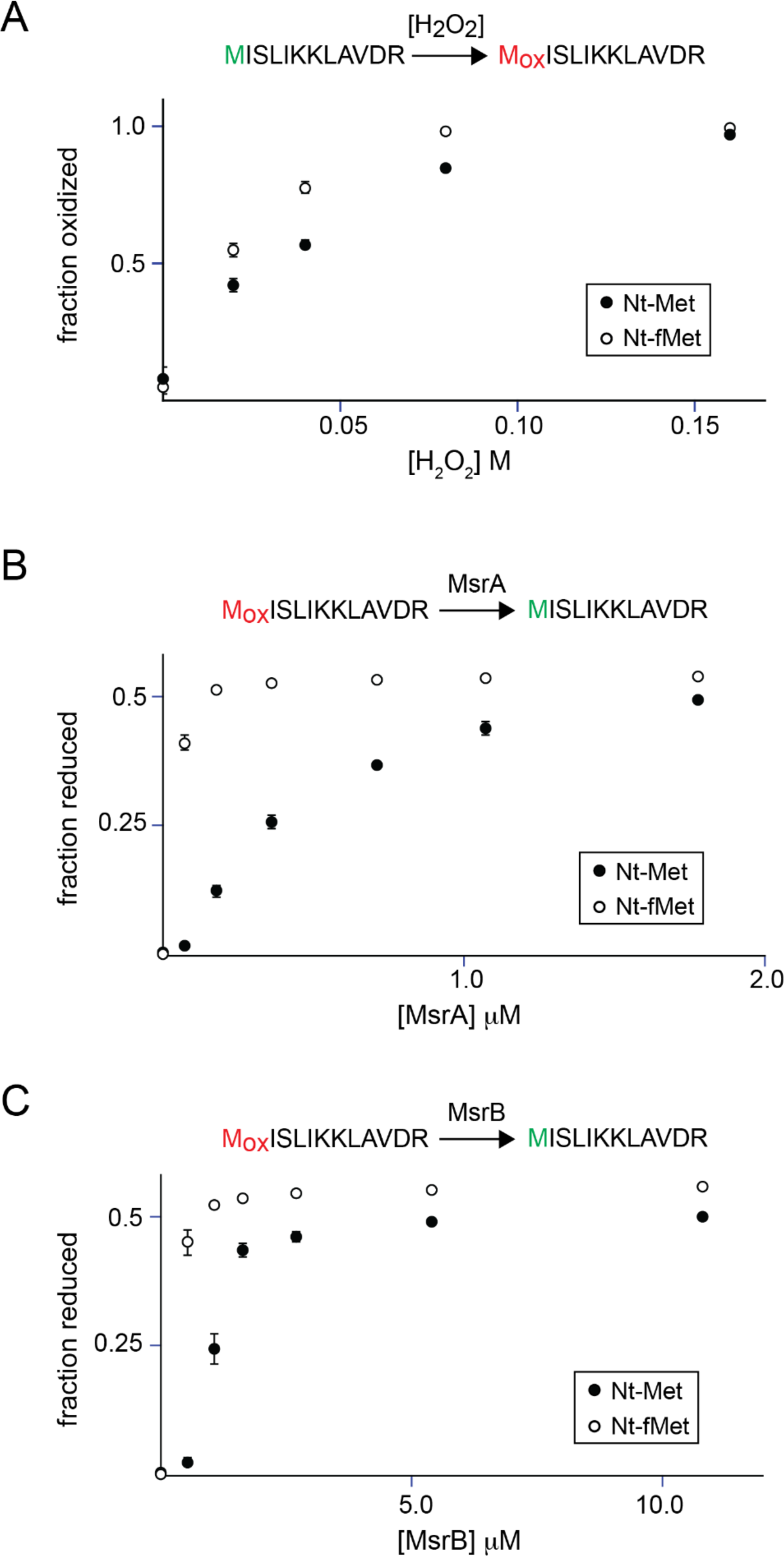
Effect of formylation on oxidation and reduction of Nt-Mets within synthetic peptides. **A)** Nt-Met oxidation as a function of H_2_O_2_ concentration. **B)** Nt-Met reduction by MsrA as a function of enzyme concentration. **C)** Nt-Met reduction by MsrB as a function of enzyme concentration. Unformylated and formylated Nt-Mets are indicated by black and white markers, respectively. Error bars represent standard deviations of three replicate experiments. See also Figure S1.

We next measured the enzymatic reduction of oxidized peptides by methionine sulfoxide reductases (Msrs) (Figure 1B,C). Nt-Met and Nt-fMet peptides were initially oxidized with hydrogen peroxide and then reduced by exposure to varying concentrations of MsrA or MsrB in presence of thioredoxin, thioredoxin reductase and NADPH. Fractional reductions of peptides were quantified by mass spectrometry (Figure S1). Since chemical oxidation is non-stereoselective and MsrA and MsrB reduce S- and R-methionine sulfoxides, respectively, complete reduction by each Msr is expected to yield an overall fractional reduction of 0.5. We observed that both MsrA and MsrB exhibited significantly enhanced reduction efficiencies for peptide substrates containing oxidized Nt-fMet relative to Nt-Met.

We determined the kinetic parameters of Msr-mediated reduction of Nt-Met- and Nt-fMet-containing substrates by measuring reduction rates as a function of peptide substrate concentrations (Table 1, Figure S1.) K_m_ of Nt-fMet was approximately 4 and 2 folds lower than Nt-Met when acted on by MsrA and MsrB, respectively. In comparison, the effect of formylation on k_cat_ values was more subtle. Overall, formylation enhanced the catalytic efficiency (k_cat_/K_m_) of MsrA and MsrB by approximate factors of 6 and 2, respectively

**Table 1.** Kinetic parameters for MsrA and MsrB mediated reduction of Nt-Met-containing synthetic peptides.

|  | MsrA |  |  | MsrB |  |  |
| --- | --- | --- | --- | --- | --- | --- |
| | $k_{\text{cat}}$ (s<sup>-1</sup>) | $K_{\text{M}}$ (μM) | $k_{\text{cat}}/K_{\text{M}}$ (M<sup>-1</sup>s<sup>-1</sup>) | $k_{\text{cat}}$ (s<sup>-1</sup>) | $K_{\text{M}}$ (μM) | $k_{\text{cat}}/K_{\text{M}}$ (M<sup>-1</sup>s<sup>-1</sup>) |
| **Mox**ISLIKKLAVDR | 0.28 ± 0.03 | 74.0 ± 22.0 | 4,500 ± 2,600 | 0.050 ± 0.005 | 23.0 ± 9.0 | 2,200 ± 1,000 |
| **fMox**ISLIKKLAVDR | 0.22 ± 0.01 | 8.6 ± 2.3 | 26,000 ± 8,000 | 0.072 ± 0.003 | 16.0 ± 2.5 | 4,400 ± 800 |

### Proteome-wide effect of formylation on Nt-Met oxidation

We next investigated the impact of Nt-formylation on methionine oxidation kinetics on a proteome-wide scale. Analyzing diverse Nt-Met-containing peptides obtained from *E. coli* protein extracts enabled us to assess the effect of formylation on oxidation of methionines in different sequence contexts. In prokaryotes, it is known that more than 90% of N-formyl groups are removed by PDF prior to the completion of translation^32^. Thus, to obtain a measurable population of Nt-fMet-containing peptides, *E. coli* bacteria were treated with the PDF inhibitor actinonin^33^ prior to protein extraction.

In initial experiments, optimal actinonin dose concentrations and treatment times were established to maximize accumulation of formylated proteins while keeping *E. coli* viable (Figure S2). Subsequently, protein extracts obtained from actinonin-treated *E. coli* were digested with trypsin and analyzed by LC-MS/MS. In all, we were able to detect more than 2,000 unique peptides containing either Nt-Mets, Nt-fMets or non-termini methionines (internal Mets). The peptide mixture was pulse oxidized by exposure to varying concentrations of unlabeled (^16^O) hydrogen peroxide and subsequently blocked by addition of excess levels of ^18^O-labeled hydrogen peroxide. As shown previously, blocking of peptides by ^18^O-labeled hydrogen peroxide prevents artifactual oxidation of methionines during the subsequent bottom-up proteomics workflow^34,35^. After blocking, fractional oxidation during the pulse phase of the experiment can be accurately quantified by determining the relative ratio of ^16^O to ^18^O-labeled peptides from the resulting mass spectra (Figure 2A).

**Figure 2.**
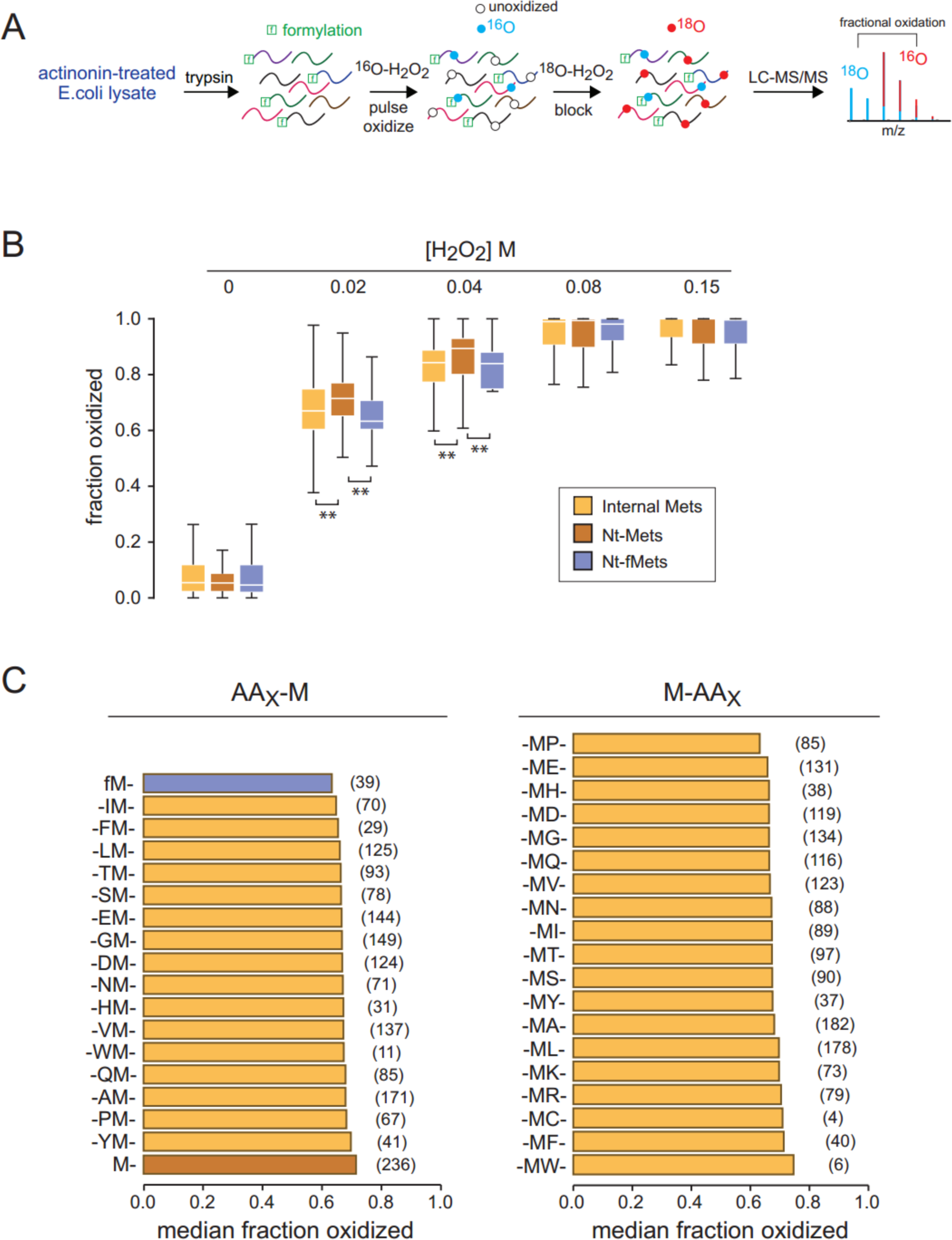
Proteome-wide analysis of the effect of formylation on Nt-Met oxidation efficiencies. **A)** Experimental design. Actinonin-treated *E. coli* were lysed and extracts were digested by trypsin. Peptides were pulse oxidized with varying concentrations of ^16^O-H_2_O_2_, quenched, and subsequently blocked with ^18^O-H_2_O_2_. Fractional oxidations of peptides were determined by measuring ^16^O/^18^O ratios using LC-MS/MS. **B)** Box plots indicating the distributions of fractional oxidation levels of peptides as a function hydrogen peroxide concentration. Peptides have been partitioned based in the presence of internal Mets (yellow), Nt-Mets (brown) and Nt-fMets (blue). Box plots indicate interquartile ranges, white lines indicate median values and whiskers represent entire ranges excluding far outliers (>2 SD). ** indicates p-value<0.01. **C)** Median fractional oxidation levels of subsets of peptides with the indicated neighboring residues on N-side (AA_X_-M) and C-side (M-AA_X_) of Mets after addition of 0.02 M H_2_O_2_. See also Figure S2, Supplementary Table S1.

Figure 2B compares the oxidation levels of peptides containing Nt-Mets, Nt-fMets and internal Mets. Consistent with previous data^36^, peptide-bound Nt-Mets were found to be more prone to oxidation than internal Mets. However, Nt-fMets had slower oxidation rates than both internal and Nt-Mets, indicating that formylation reduces the oxidation rates of methionines (Figure 2B, Supplementary Table S1).

We next used the proteomic data to investigate the influence of neighboring residues on the oxidation kinetics of peptide-bound methionines. Figure 2C compares the average oxidation levels of peptides with varying N- and C-neighboring residues. We note that this analysis was not exhaustive of all possible sequence contexts (see Methods). For example, lysine and arginine residues could not be detected at the N-side of methionines because of proteolytic selectivity of trypsin digestion. Consistent with the analysis shown in Figure 2B, the results indicated that Nt-Mets are more prone to oxidation than internal Mets regardless of the N-bound neighboring residue, and that formylation attenuates the oxidation of Nt-Mets. However, proteome-wide differences in oxidation levels of Nt-Mets, Nt-fMets and internal Mets were relatively slight (median oxidation levels were within ∼15% of each other) and variable between peptides. For example, although formylation decreased oxidation levels of Nt-Mets for most peptides in our proteome-wide experiment, within the synthetic peptide analyzed in Figure 1A, it in fact increased methionine oxidation levels slightly. We therefore conclude that formylation has a modest and variable effect on oxidation of N-terminal methionines.

### Proteome-wide effect of formylation on Msr-mediated reduction of Nt-Mets

We next analyzed the impact of Nt-formylation on methionine reduction kinetics using bottom-up proteomics (Figure 3A, S3, Supplementary Table S2). To generate a diverse mixture of oxidized substrates containing formylated and unformylated peptides, actinonin-treated *E. coli* extracts were trypsinized and oxidized with ^16^O-H_2_O_2_. The peptides were then enzymatically reduced with MsrA, MsrB or both for varying times in the presence of thioredoxin, thioredoxin reductase and NADPH. Reactions were subsequently quenched and peptides were blocked with ^18^O H_2_O_2_. ^16^O/^18^O ratio of peptides were quantified by LC-MS/MS as before.

**Figure 3.**
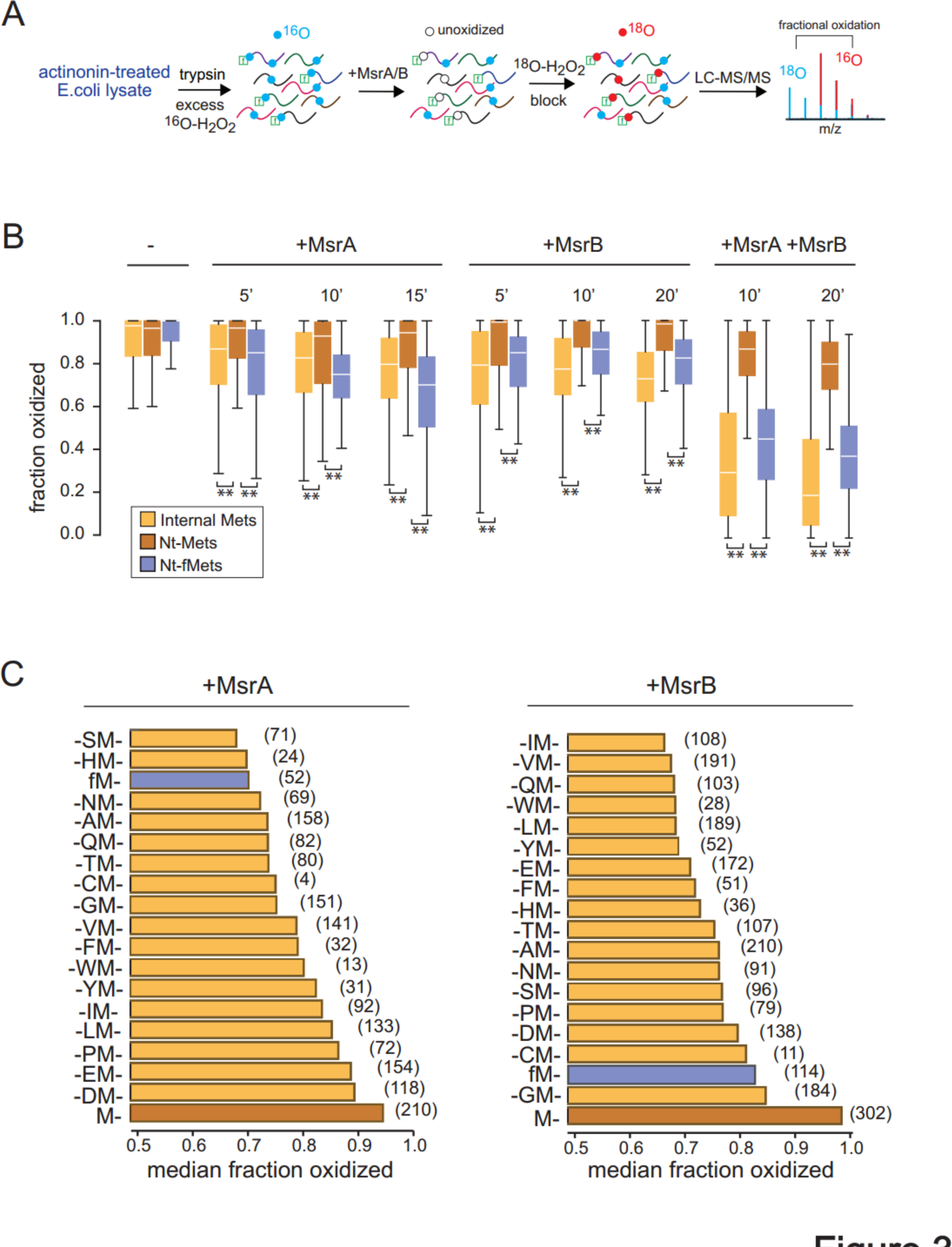
Proteome-wide analysis of the effect of formylation on Nt-Met reduction efficiencies by MsrA and MsrB. **A)** Experimental design. Actinonin-treated *E. coli* were lysed and extracts were digested by trypsin. Peptides were fully oxidized with ^16^O-H_2_O_2_ and subsequently reduced with MsrA or MsrB or both for variable times. Peptides were subsequently blocked with ^18^O-H_2_O_2_ and fractional oxidation levels were determined by measuring ^16^O/^18^O ratios using LC-MS/MS. **B)** Box plots indicating distributions of fractional oxidation levels of peptides as described in Figure 2B. **C)** Median fractional oxidation levels of subsets of peptides with the indicated neighboring residues on N-side (AA_X_-M) of Mets after exposure to MsrA or MsrB for 10 minutes. See also Figure S3, Supplementary Table S2.

Consistent with our observations in synthetic peptides, Nt-fMet residues exhibited rapid reduction when subjected to MsrA, MsrB or both, while Nt-Met residues had comparatively slower reduction rates (Figure 3B). Further sequence analysis of peptide substrates indicated that MsrA and MsrB have distinct sequence selectivities (Figure 3C). For example, MsrA was most efficient at reducing methionine sulfoxides that were followed by polar residues such as serine (S), asparagine (N), and glutamine (Q) while MsrB was most efficient at reducing methionine sulfoxides followed by hydrophobic residues such as isoleucine (I), valine (V), tryptophan (W) and tyrosine (Y). Proline residues downstream of methionine sulfoxides significantly reduced the efficiency of both enzymes (Figure S3). However, among all possible neighboring residue contexts for methionines, Nt-Mets were the least efficient substrates for both MsrA and MsrB. Formylation of oxidized Nt-Mets greatly enhanced their reduction by both enzymes.

### Proteome-wide effect of acetylation on Msr-mediated reduction of Nt-Mets

We next sought to investigate the effects of α-amine modifications other than formylation on Msr-mediated reduction of methionines. N-terminal acetylation is a prevalent co- and post-translational protein modification, particularly in eukaryotes^26^. To determine the impact of acetylation on Msr-catalyzed reduction of Nt-Mets, we conducted proteome-wide reduction experiments on chemically acetylated *E. coli* extracts. Trypsinized extracts were treated with acetic anhydride to generate N-term acetylated peptides (Figure S4A) and subsequently mixed with untreated extracts to produce a mixture of peptides containing both acetylated and unacetylated Nt-Mets. As before, the peptide mixture was oxidized, subsequently reduced with MsrA, MsrB or both, and levels of oxidation quantified by LC-MS/MS (Figure 4A, Supplementary Table S3). The results indicated that acetylated Nt-Mets (Nt-AcMets) have faster rates of reduction than unmodified Nt-Mets (Figure 4B). Thus, we conclude that both formylation and acetylation enhance the reduction efficiency of Nt-Mets by MsrA and MsrB.

**Figure 4.**
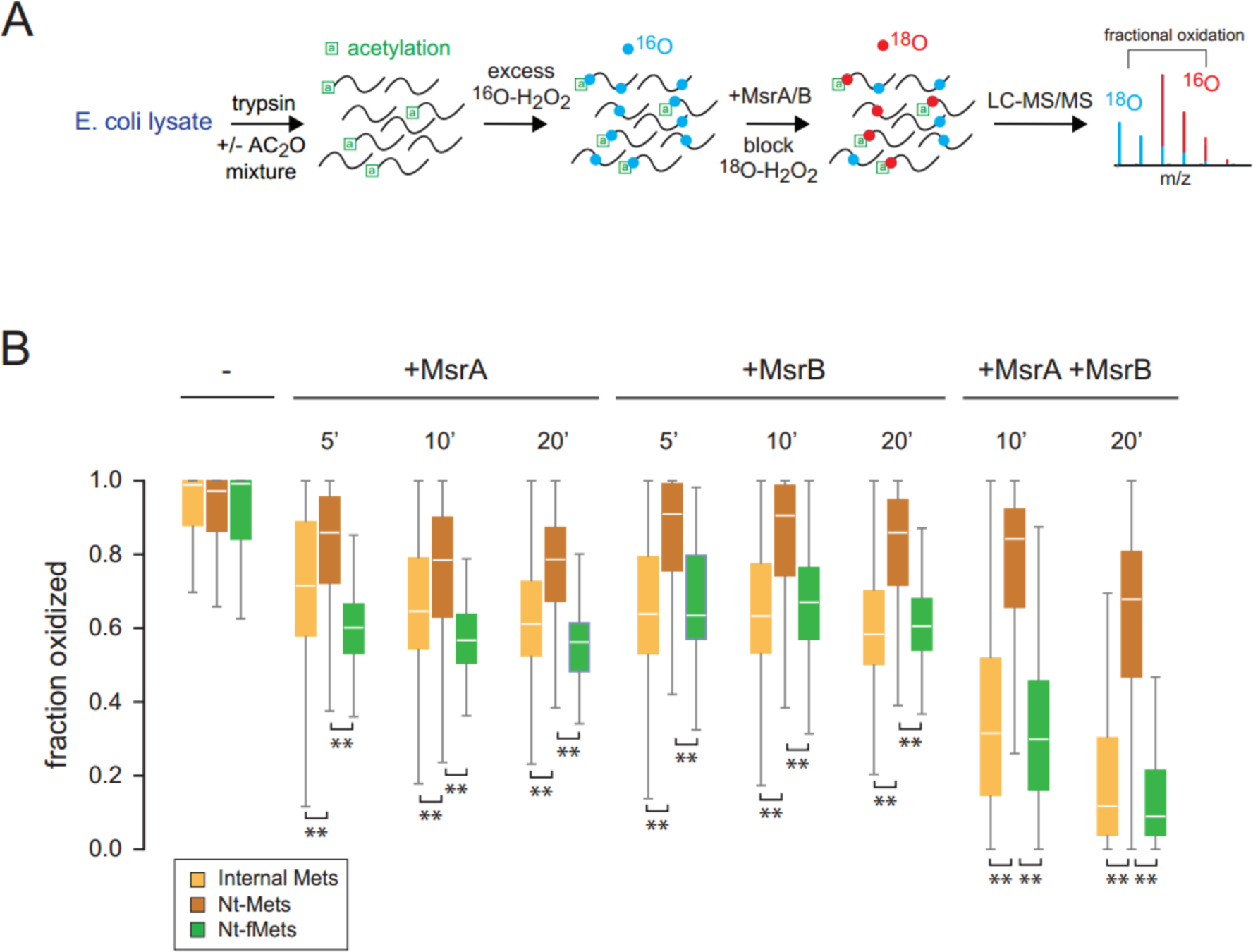
Proteome-wide analysis of the effect of acetylation on Nt-Met reduction efficiencies by MsrA and MsrB. **A)** Experimental design. *E. coli*-derived peptides were treated with acidic anhydride and mixed with untreated trypsinized peptides at 1:1 ratios. Peptides were fully oxidized with ^16^O-H_2_O_2_ and subsequently reduced with MsrA or MsrB or both for variable times. Peptides were subsequently blocked with ^18^O H_2_O_2_ and fractional oxidation levels were determined by measuring ^16^O/^18^O ratios using LC-MS/MS. **B)** Box plots indicating distributions of fractional oxidation levels of peptides as described in Figure 2B. See also Figure S4, Supplementary Table S3.

### Oxidation of Nt-Mets prevents their co-translational processing and Msrs rescue this inhibition

Co-translational processing of Nt-fMets encompasses their deformylation by PDF and, in some cases, their subsequent excision by MAP^37,38^. In bacteria, the ability to properly process Nt-Mets is essential for viability as loss of PDF and MAP activities are known to be lethal^39,40^. We reasoned that oxidation of Nt-Mets in nascent polypeptides may prevent their subsequent processing by PDF and MAP. Thus, efficient Msr-mediated reduction of oxidized Nt-fMets may provide a mechanism for rescuing the deleterious inhibition of their subsequent processing. To investigate this possible effect, we analyzed the impact of Nt-fMet oxidation on *in vitro* activities of PDF and MAP using synthetic peptides as substrates. The data indicated that Nt-fMet oxidation almost fully inhibits its deformylation by PDF, and Nt-Met oxidation almost fully inhibits its hydrolysis by MAP (Figure 5A,B, Figure S5A,B). Accordingly, tandem processing of Nt-fMets by combined addition of PDF and MAP were inhibited by oxidation (Figure 5C, Figure S5C). As predicted, the inclusion of MsrA and MsrB in the assay rescued this inhibition and enabled the proper processing of polypeptides that initially harbored oxidized Nt-fMets. Hence, the ability of Msrs to efficiently reduce formylated methionine sulfoxides enhances their capacity to revert the inhibitory effects of oxidation on Nt-Met processing *in vitro*.

**Figure 5.**
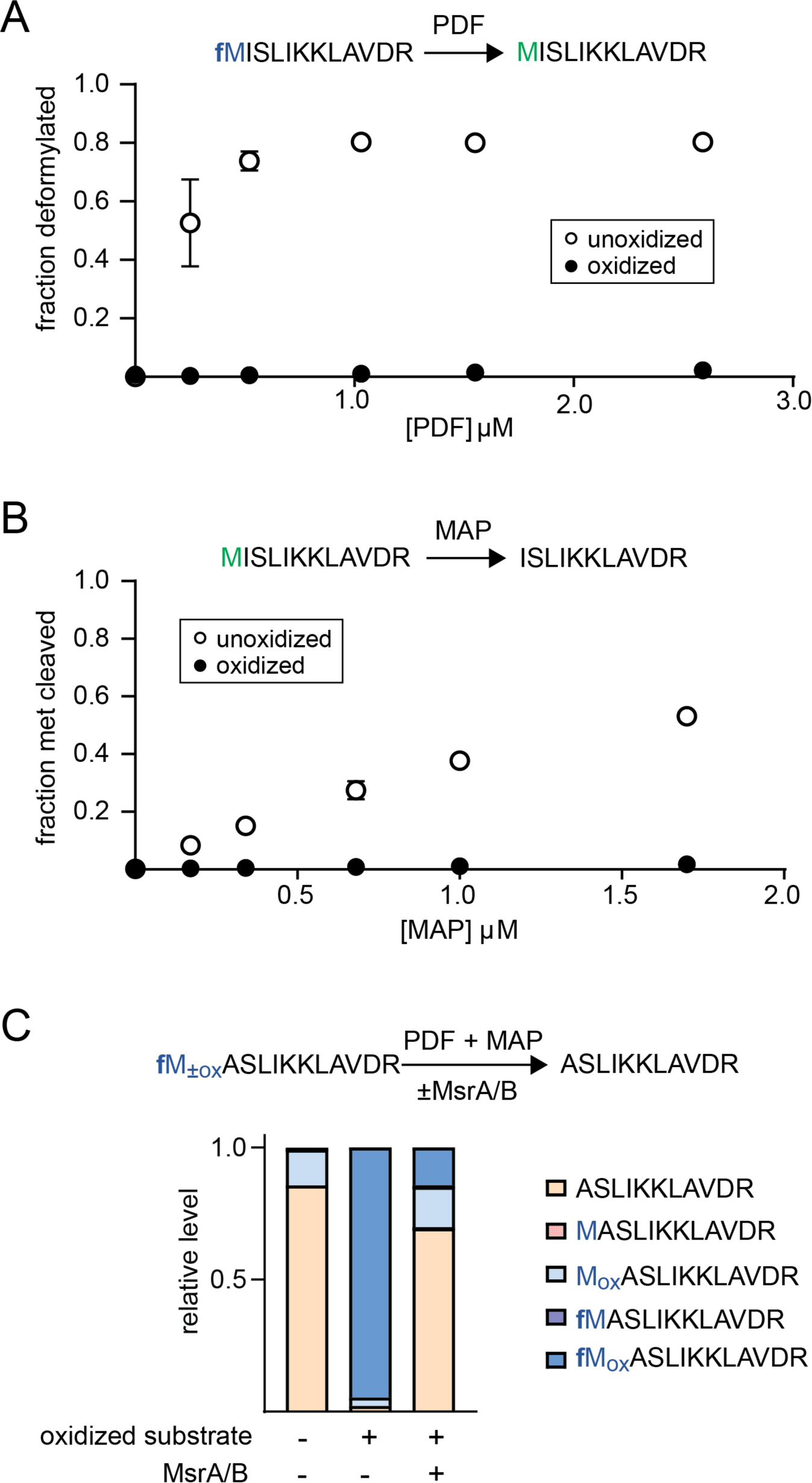
Effect of oxidation on the deformylation and cleavage of Nt-Mets within synthetic peptides. **A)** Nt-fMet deformylation as a function of PDF concentration. **B)** Nt-Met cleavage as a function of MAP concentration. In (A) and (B), unoxidized and oxidized Nt-Mets are indicated by black and white markers, respectively. Error bars represent standard deviations of three replicate experiments. **C)** Nt-fMet deformylation and subsequent cleavage by combined addition of PDF and MAP in the presence or absence of MsrA and MsrB. The bar plot indicates the relative proportion of each of the possible resulting modified forms of the peptide substrate at the end of the experiment. See also Figure S5.

### Lack of formylation escalates levels of oxidized Nt-Mets in vivo

The above *in vitro* data suggest that formylation of Nt-Mets may curtail their oxidation. To assess this hypothesis *in vivo*, we generated *E. coli* deficient in Nt-Met formylation by knocking out the *fmt* gene (*Δfmt*) and investigated their response to oxidative stress. Consistent with previous reports^20^, the *Δfmt E. coli* strain was viable but exhibited significantly slower growth rates relative to wildtype. To compare methionine oxidation levels between *Δfmt* and wildtype cells, bacteria were incubated in the presence or absence of 20 mM hydrogen peroxide for 1 hour and their extracts were trypsinized and analyzed by LC-MS/MS. We expected endogenous methionine oxidation levels to be significantly lower *in vivo* in comparison to the *in vitro* assays above (where peptides were chemically oxidized), thus making the exact quantitation of oxidation stoichiometry by ^18^O-H_2_O_2_ treatment difficult^34^. Therefore, we instead compared numbers of peptide spectral matches (PSMs) for different categories of unoxidized and oxidized peptides between wildtype and *Δfmt* cells (Figure 6, Supplementary Table S4). Overall, we were able to detect ∼35,000 total PSMs for each sample. As expected, higher numbers of oxidized methionine-containing peptides were detected in hydrogen peroxide treated cells compared to untreated controls. However, relative to wildtype, numbers of oxidized methionine-containing peptides were higher in the *Δfmt* strain in both untreated and hydrogen peroxide treated cultures. This difference was particularly evident in peptides mapped to N-termini of proteins that harbored initiator methionines. Additionally, numbers of PSMs corresponding to N-terminal peptides with unhydrolyzed initiator methionines were greater in hydrogen peroxide treated and *Δfmt* cells. Together, the data indicate that formylation facilitates the reduction of oxidized Nt-Mets *in vivo* and promotes their proper processing.

**Figure 6.**
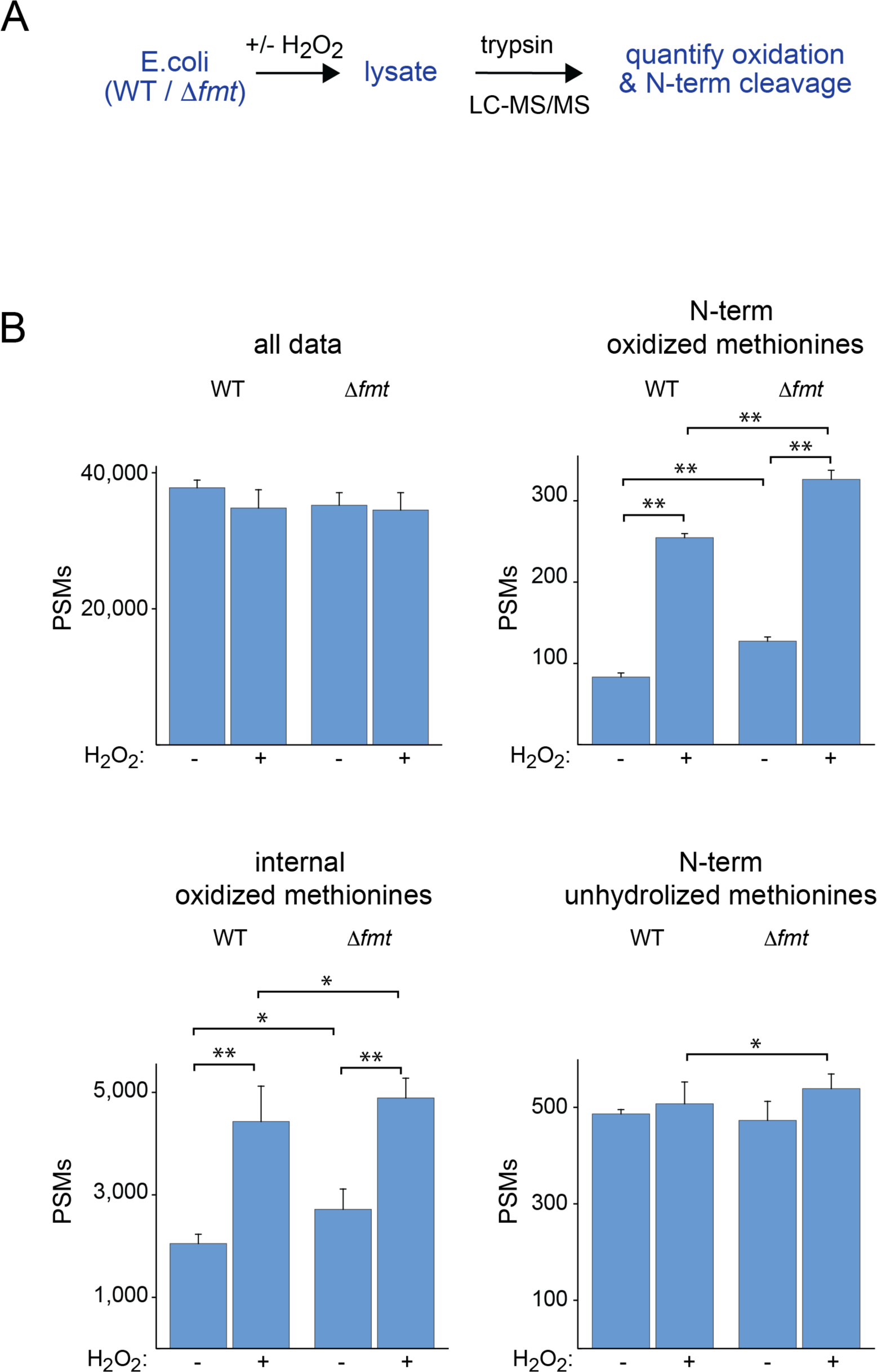
Methionine oxidation levels in *τιfmt E. coli*. **A)** Experimental design. Wildtype and *τιfmt E. coli* were grown under normal conditions or exposed to oxidative stress in the form of hydrogen peroxide. Extracts were collected, trypsinized and analyzed by LC/MS-MS. **B)** Total number of detected PSMs, PSMs with oxidized Nt-Mets, PSMs with oxidized internal Mets and PSMs with uncleaved Nt-Mets. Error bars represent standard deviations of three replicate experiments. ** indicates p-value<0.01 and * indicates p-value<0.05. See also Supplementary Table S4.

## Discussion

We have shown that within polypeptides, Nt-Mets are particularly prone to oxidation and are inefficiently reduced by MsrA and MsrB. Formylation and acetylation of Nt-Mets alter their redox properties and greatly enhance their reduction by Msrs. Proteome-wide analyses indicated that this effect is consistent across diverse peptide sequences and largely driven by alterations in the Km of the enzymatic reactions.

It is known that MsrA and MsrB are more efficient at binding and reducing protein-bound methionine sulfoxides in comparison to free methionine sulfoxides^41,42^. The selectivity of Msrs for protein-bound substrates suggests that their active sites either favorably interact with peptide amide bonds flanking protein-bound methionine sulfoxides, or unfavorably interact with charged amino and/or carboxyl groups of free methionine sulfoxides^43–45^. However, to date, experimentally determined substrate-bound structures of Msrs have not been able to provide a definitive structural basis for this selectivity^46^. Our results are consistent with these previous observations and suggest that the formylation and acetylation of Nt-Mets may enhance their Msr-mediated reduction either by removing the positive charge of the N-terminal amino group that unfavorably interacts with the enzyme, or by generating an amide group that favorably interacts with the enzyme. However, the structural details of these interactions remain to be determined.

It is well known that methionine formylation plays an important role in protein synthesis within prokaryotic cells^47^. MTF catalyzes the formylation of methionines bound to initiator tRNAs that are subsequently incorporated as the first residue of nascent polypeptides during translational initiation^20,47^. The formyl group of fMet-tRNA_i_^Met^ promotes its interaction with initiation factor 2 (IF2) and enhances the efficacy of translational initiation^48^. However, the formylation of Nt-Met is not strictly essential for protein synthesis or cell viability, and for many bacteria, the deletion of the fmt gene is not lethal. Alternatively, it has been suggested that formylated Nt-Mets can act as degrons and promote the proteolysis of misfolded proteins following translation^28^.

This study highlights a third possible consequence of methionine formylation: prevention of methionine oxidation during the course translation. Unlike formylation, the deformylation of Nt-Met in nascent polypeptides is essential for bacterial viability and the deletion of the PDF-gene (*def*) is known to be lethal^39^. The presence of a formyl group on Nt-Mets inhibits their subsequent cleavage by MAP, which is required for the activity a number of essential enzymes^23^. In this study, we have shown that methionine oxidation inhibits both deformylation by PDF and cleavage by MAP. Thus, formylation expedites the reduction of oxidized Nt-Mets Msrs and enables their subsequent processing.

It is also possible that the rapid oxidation and reduction of Nt-fMets acts as a scavenger mechanism, depleting local concentrations of reactive oxygen species (ROS) and protecting internal residues in nascent polypeptides from oxidation. Similar scavenging functions have been attributed to reversible methionine oxidation within the cell in different biological contexts^15,49–51^. As formyl groups within most nascent proteins are removed by PDF early during the course of translational elongation, they are unlikely to have a significant impact on the oxidation of most internal residues. Consistent with this idea, our data indicated that the ablation of formylation in *E.coli* primarily impacted the oxidation of Nt-Mets and not internal Mets.

Whether the cellular process of formylation has evolved to facilitate the reduction of Nt-Mets, or conversely, Msr function has evolved to accommodate and harness Nt-Met formylation, cannot be determined by this study. It is also possible that the efficiency of Msrs in reducing Nt-fMets is simply a fortuitous consequence of their inherent selectivity for protein-bound methionines that are flanked by peptide amide bonds. Regardless of its evolutionary origins, the synergy between formylation and Msr-mediated reduction appears to reduce levels of oxidized Nt-Mets and promote their subsequent processing.

## Methods

### Enzyme expression and purification

*E. coli* methionine sulfoxide reductase A (MsrA), methionine sulfoxide reductase B (MsrB), thioredoxin (TrxA), thioredoxin reductase (TrxB), peptide deformylase (PDF) and methionine aminopeptidase (MAP) were cloned in pET151/D-TOPO vectors (Thermo) downstream of T7 promoter and N-terminally tagged with 6x His-tag. The plasmids were transformed into BL21/DE3 competent cells (Thermo) and protein expression was induced by 400 µM IPTG (American Bio) overnight at 25 ℃. Protein purification was conducted with HisPur Ni-NTA resin (Thermo Scientific) on a Poly-Prep Chromatography Column (BioRad). Imidazole (Sigma) was added at concentrations ranging from 150 mM to 250 mM to elute protein from the resin. Eluted proteins were buffer exchanged using dialysis (dialysis cassette, 7K MWCO, 3ml, Thermo Fisher). The terminal buffers were 50 mM Tris, pH 7.5 for MsrA, MsrB, TrxA and TrxB; 50 mM HEPES, 0.2 mM CoCl_2_, pH 7.4 for PDF; and 100 mM K_2_SO_4_, 0.2 mM CoCl_2_, pH 7.4 for MAP. Purified proteins were stored at -80 °C in 10% glycerol.

### Oxidation and reduction of synthetic peptides

All peptides in their Nt-Met and Nt-fMet forms were synthesized by GenScript. For oxidation experiments, Nt-Met and Nt-fMet forms of peptides were resuspended in 10% ACN and mixed at 1:1 ratio. Oxidation was performed in 0 , 20, 40, 80 and 150 mM of H_2_O_2_ in water for 20 minutes at 37 ℃ at peptide concentrations of 0.18 mM. Three replicates of each experiment were conducted. Oxidation was quenched by addition of 300 mM sodium sulfide (Sigma). Peptides were desalted using homemade C18 columns and eluted with 0.1% formic acid in 50/50 formic acid/acetonitrile to a final concentration of 15 µg/mL.

For reduction experiments, Nt-Met and Nt-fMet forms of peptides were resuspended in water and mixed at 1:1 ratio. The mixture was fully oxidized with 150 mM H_2_O_2_ at 37 ℃ for 30 minutes. Excess H_2_O_2_ was removed by lyophilization. Oxidized peptides were resuspended in 50 mM Tris (pH 7.5) at a concentration of 1mg/mL. Reduction reactions were carried out by addition of 0.07, 0.18, 0.36, 0.71, 1.07 and 1.78 µM of MsrA, or 0.54, 1.08, 1.62, 2.69, 5.39 and 10.79 µM of MsrB in presence of 7 µM thioredoxin, 5 µM thioredoxin reductase and 10 mM NAPDH in 50 mM Tris buffer (pH 7.5) in a total volume of 60 µL. The peptide concentration was 0.18 mM. Three replicates of each experiment were conducted. The reduction reaction was carried out for 10 minutes at 37℃ and quenched by addition of 60 µL acetonitrile (ACN). Peptides were lyophilized and resuspended in 50 µL 0.1% formic acid and desalted using homemade C18 columns and eluted with 0.1% formic acid in 50/50 formic acid/ACN to a final concentration of 15 µg/mL. The samples were analyzed in mass spectrometer.

### Michaelis-Menten kinetics

Kinetic analyses of reduction reactions were carried out by monitoring the oxidation of NADPH through loss of absorbance at 340 nm, as described previously^9^. Nt-Met and Nt-fMet peptides were oxidized as above and reconstituted in 50 mM Tris-HCl (pH 7.5) at a concentration of 5 µg/µL. Reduction reactions were carried out in five replicate experiments by addition of 0.75 µM MsrA (Nt-Met), 0.25 µM MsrA (Nt-fMet), or 1 µM MsrB (Nt-Met and Nt-fMet). Thioredoxin and Thioredoxin reductase were added at 10x Msr concentration. NADPH concentrations were either 64 µM (replicate 1 and 2) or 200 µM (replicate 3, 4 and 5). The buffer was 50 mM Tris-HCl (pH 7.5). Oxidized Nt-Met and Nt-fMet peptide substrate concentrations were 0, 10, 20 40, 80, 160, 320 and 640 µM. A NADPH control at 10 µM substrate was also included as an absorbance baseline.

Reactions were carried out in 25 µL volumes in UV-Star 96-well microplates (Grenier Bio-one). Absorbance at 340 nm was recorded using a SpectraMax M2e plate reader at 37°C every 10 seconds for 10 minutes (replicate 1 and 2) or 20 minutes (replicate 3, 4 and 5). Data analysis was carried out using Mathematica (Wolfram). Initial velocities were measured by linear regression and corrected for background NADPH decay by subtracting the rate of the 10 µM substrate control. The peptide substrate concentrations used for data analyses were assumed to be half of the experimental concentrations of the racemic peptide mixtures, as MsrA and MsrB are known to be stereoselective for the S-and R-MetO diastereomers, respectively. The initial peptide substrate concentrations were also corrected for incomplete oxidation as measured by mass spectrometry. The resulting initial velocity versus substrate plot was fit to the Michaelis-Menten equation by non-linear regression to measure K_M_ and k_cat_.

### Generation of peptide samples for proteomic experiments

For peptide samples containing formylated Nt-Mets, wildtype K12 *E.coli* were treated with actinonin (Sigma) at a final concentration of 60 µg/mL when OD_600_ reached 1.0 and incubated for 0, 10, 30, 60 and 120 minutes at 37 ℃. Cell pellets were resuspended in lysis buffer containing 5% SDS in 50 mM TEAB. The cell suspension was lysed by sonication at 30 amp for 5 minutes and clarified of cell debris by centrifugation at 16000 xg for 5 min. Following lysis, protein concentration was quantified by bicinchoninic acid (BCA) assay (Thermo). 25 µg aliquots of protein were diluted to a final concentration of 1 mg/mL. Disulfide bonds were reduced by adding 2 mM dithiothreitol (DTT) (Sigma) and incubated at 55 ℃ for 1 hour. Subsequently, alkylation was performed with 10 mM iodoacetamide (IAA) (Sigma) for 30 min at room temperature in the dark. Samples were acidified by 1.2% phosphoric acid and diluted 7-fold with 90% methanol in 100 mM TEAB. Diluted samples were transferred to S-trap columns (Protofi) by centrifugation at 4000 xg for 1 min. Columns were washed twice by 90% methanol in 100 mM TEAB. Trypsin (Pierce) was added to the S-trap column at a ratio of 1:25 (trypsin/protein) and the digest reaction was allowed to continue overnight at 37 ℃. Peptides were eluted with 40 µL of 0.1% trifluoroacetic acid (TFA)in water (Pierce), followed by 40 µL of 0.1% TFA in 50/50 ACN/water. Digested peptides were lyophilized and resuspended in 0.1% TFA to 0.5 mg/mL prior to mass spectrometry analysis.

For peptide samples containing acetylated Nt-Mets, 25 µg of actinonin-untreated *E.coli* protein extracts were digested into peptides as described above. 0.2% acetic anhydride in 1M ammonium bicarbonate was added to the peptides at 1:4 ratio (v/v, peptide/acetic anhydride) and incubated 30 minutes at 37 ℃. The acetylated peptides were diluted in a 1:1 ratio with 0.1% TFA in water and desalted in the homemade C18 columns prior to mass spectrometry analysis.

### Proteome-wide oxidation kinetics

Digested peptides prepared as described above were resuspended in 0.1% TFA in water to a final concentration of 0.3 mg/mL. Oxidation was performed in 0, 20, 40, 80 and 150 mM of ^18^O H_2_O_2_ in water (Sigma) and incubated for 30 minutes at 37 ℃. Oxidation was quenched by 300 mM sodium sulfide (Sigma) and desalted with homemade C18 columns. To block unoxidized methionines, peptides were lyophilized and resuspended in 150 mM ^16^O H_2_O_2_ in water and incubated for 30 min at 37 ℃. The remaining hydrogen peroxide was removed through lyophilization. Samples were resuspended in 0.1% TFA to 0.5 mg/ml and analyzed mass spectrometry as described below.

### Proteome-wide reduction kinetics

Digested peptides prepared as described above were resuspended in 0.1% TFA in water to a final concentration of 0.3 mg/mL. Peptides were fully oxidized by incubation in 150 mM of ^16^O H_2_O_2_ for 30 minutes at 37 ℃. Excess hydrogen peroxide was removed by lyophilization and oxidized peptides were resuspended in 50 mM Tris (pH 7.5) and 50 mM DTT. The reduction reaction was performed in presence of 0.54 µM MsrA or 2.02 µM MsrB or both for 5, 10 or 20 minutes at 37 ℃. The reaction was subsequently quenched by ACN. The samples were subsequently diluted with Tris buffer to reduce acetonitrile concentration below 20% and enzymes were removed by 3K-cutoff centrifugal filters (Amicon). To block unoxidized methionines, peptides were lyophilized and resuspended in 150 mM ^18^O H_2_O_2_. for 30 min at 37 ℃. Samples were desalted with C18 column, lyophilized,resuspended in 0.1% TFA in water, and analyzed by mass spectrometry as described below.

### Deformylation and excision of Nt-Met in the presence and absence of Msrs

Deformylation and excision assays were conducted by addition of PDF or MAP, respectively, to either fully oxidized or unoxidized peptides. Peptides were resuspended in 100 mM KH_2_PO_4_ 0.2 mM CoCl_2_, pH 7.4 at a concentration of 1 mg/mL. 0.2 mM of each peptide was incubated with 0.25, 0.52, 1.03, 1.55 and 2.59 µM of PDF or 0.17, 0.34, 0.68, 1.0 and 1.7 µM of MAP in 50 µL volumes. Three replicate experiments were conducted for each reaction. For Msr rescue experiments, reactions were performed in the presence of 1.03 µM PDF, 0.68 µM MAP, 7 µM thioredoxin, 5 µM thioredoxin reductase, 6 mM NADPH, 0.71 µM MsrA and 2.69 µM MsrB. After 10 minutes of incubation at 37 ℃, reactions were quenched by addition of 50 µL ACN. Enzymes and salts were subsequently removed using C18 desalting columns. Peptides were lyophilized and resuspended in 50/50 0.1% formic acid/ACN at 15 µg/mL concentrations and analyzed by mass spectrometry as described below.

### Generation, growth and treatment of Δfmt E. coli

Deletion of *fmt* in K-12 W3110 *E. coli* was conducted using the method of Datsenko and Wanner^52,53^. Briefly, a KanR cassette including flippase recognition targe (FRT) sites was amplified from the pKD13 vector by PCR, digested by DpnI, and purified using a QIAquick Gel Extraction Kit (Qiagen). The forward PCR primer (5’-GAAAAACTGGATCGTCTGAAAGCCCGGGCTTAAGGATAAGAACTAACGTGATTCC GGGGATCCGTCGACC-3’) included a standard 20 nt region homologous to the upstream portion of the KanR cassette and was tailed by 50 nts of homology including the start codon of *fmt* and the region immediately upstream. The reverse primer (5’-GGCAAGACCGGGCTTAGAAGAGTGGACTATCAGACCAGACGGTTGCCCGGTGTAGGCTGGAGCTGCTTCG-3’) included a standard 20 nt region homologous to the downstream portion of the KanR cassette and was tailed by 50 nts of homology to the final 21 nts of *fmt* including the stop codon, along with the region immediately downstream. *E. coli* carrying the Red recombinase expression vector pKD46 (encoding the λ phage genes, gam, exo and beta under an araB promoter) were then transformed with the above PCR product by electroporation and transformants were selected by kanamycin. To remove the pkD46 vector, the cells were incubated at 42°C for 3 hours and curing of the plasmid was confirmed by an inability to grow on ampicillin. To verify knockouts, colony PCR was performed on transformants using primers upstream (5’-CCGCGACGGTAAACCATTTG-3’) and downstream (5’-GCTCGACGACTTGTTCAACG-3’) of the *fmt* locus. The knockout was verified by the expected change in the size of the PCR product as well as through Sanger sequencing using the same primers used for the colony PCR.

*Δfmt* and W3110 WT bacteria were grown to an OD_600_ ∼ 0.5-0.8 in an overnight culture. 2 mL of the bacteria was diluted to an OD_600_ of 0.5 and incubated with 0 mM or 20 mM hydrogen peroxide for 1 hour at 37 ℃, shaking at 200 rpm. The bacteria were pelleted by centrifugation at 8,000 xg for 10 minutes. Pellets were lysed in 5% SDS by sonification. Proteins were digested following the S-trap and tryptic digestion protocol described above. Three biological replicates were analyzed.

### LC-MS/MS analysis

For analysis of synthetic peptides, samples were reconstituted in 50% acetonitrile, 0.1% formic acid at a concentration of 10 µg/mL and transferred to an autosampler vial. A pseudo-infusion was performed by injecting 50 µL using a Dionex Ultimate 3000 connected to a Q Exactive Plus mass spectrometer (Thermo Fisher) without the use of a column. The bolus of sample was introduced to the mass spectrometer using a flow rate of 100 µL/min. Solvent A was 0.1% formic acid in water, and solvent B was 0.1% formic acid in acetonitrile, each at 50% during the length of the 3-minute method. Peptides were ionized using a HESI source in positive mode. Full scans were performed with a resolution of 70,000 at m/z 200, with an AGC target of 1e6 and a maximum injection time of 240 ms over a scan range of 300-2000 m/z.

For proteome-wide analysis of *E. coli* peptide extracts, samples were injected onto a homemade 30 cm C18 column with 1.8 um beads (Sepax), with an Easy nLC-1200 HPLC (Thermo Fisher), connected to a Fusion Lumos Tribrid mass spectrometer (Thermo Fisher). Solvent A was 0.1% formic acid in water, while solvent B was 0.1% formic acid in 80% acetonitrile. Ions were introduced to the mass spectrometer using a Nanospray Flex source operating at 2 kV. The gradient began at 3% B and held for 2 minutes, increased to 10% B over 5 minutes, increased to 38% B over 38 minutes, then ramped up to 90% B in 3 minutes and was held for 3 minutes, before returning to starting conditions in 2 minutes and re-equilibrating for 7 minutes, for a total run time of 60 minutes. The Fusion Lumos was operated in data-dependent mode, with MS1 scans acquired in the Orbitrap, and MS2 scans acquired in the ion trap. The cycle time was set to 1.5 seconds. Monoisotopic Precursor Selection (MIPS) was set to Peptide. The full scan was done over a range of 375-1400 m/z, with a resolution of 120,000 at m/z of 200, an AGC target of 4e5, and a maximum injection time of 50 ms. Peptides with a charge state between 2-5 were picked for fragmentation. Precursor ions were fragmented by collision-induced dissociation (CID) using a collision energy of 30% with an isolation width of 1.1 m/z. The Ion Trap Scan Rate was set to Rapid, with a maximum injection time of 200 ms, an AGC target of 1e4. Dynamic exclusion was set to 20 seconds.

### Proteomic data analysis and measurement of ^16^O/^18^O peptide ratios

Raw files for all samples were searched against the Escherichia coli K12 UniProt database (UP000000625_83333, downloaded 4/3/2022) using Fragpipe software^54^. Peptide and protein quantifications were performed with Fragpipe default parameter settings. ^18^O, ^16^O, and N-terminal formylation were set as variable modifications; carbamidomethyl cysteine was set as a fixed modification. Raw files were converted to .ms1 format using MSConvert software^55^. The vendor-supplied peak picking algorithm and the threshold peak filter set to the top 2000 peaks in each scan.

Fragpipe-supplied PSM files were used for quantitation of PSM numbers. Quantification of ^16^O/^18^O labeled methionine ratios was carried out as previously described^34,35^. Briefly, MS1 spectra corresponding to specific methionine-containing peptides were extracted from the data. These spectra typically contain overlapping isotopic envelopes corresponding to ^16^O- and ^18^O-labeled forms of peptides. The population of each form was measured by deconvoluting and measuring the relative intensities of the two isotopic envelopes using a previously described algorithm^34,35^.

### Author Contributions

RT & SG conceived of the experiments. RT, MH, PB, SB, KC, KS, KW & JH performed the experiments. RT, MH, PB, KW & SG analyzed the data. RT & SG wrote the initial manuscript.

### Funding Sources

This work was supported by grants from the National Institutes of Health (R35 GM119502 and S10 OD025242 to SG).

### Notes

The authors declare no competing financial interest.

## Acknowledgements

We thank the members of the Ghaemmaghami lab at the University of Rochester for helpful discussions and suggestions.

## Data Availability

All raw and processed data are available in the included Supporting Information and at the ProteomeXchange Consortium via the PRIDE partner repository (accession number PXD048776). Currently, the data can be accessed with the username and password **YS9ELiGe**.

## Abbreviations

LC-MS/MS: liquid chromatography-tandem mass spectrometry
SPROX: Stability of Proteins from Rates of Oxidation
MObB: Methionine Oxidation by Blocking
MTF: methionyl-tRNA formyltransferase
PDF: peptide deformylase
MAP: methionine aminopeptidase
MsrA: methionine sulfoxide reductase A
MsrB: methionine sulfoxide reductase B
DTT: dithiothreitol
IAA: iodoacetamide
TFA: trifluoroacetic acid
ACN: acetonitrile
PSM: peptide-spectrum match

## Supplementary Figures

**Figure S1.**
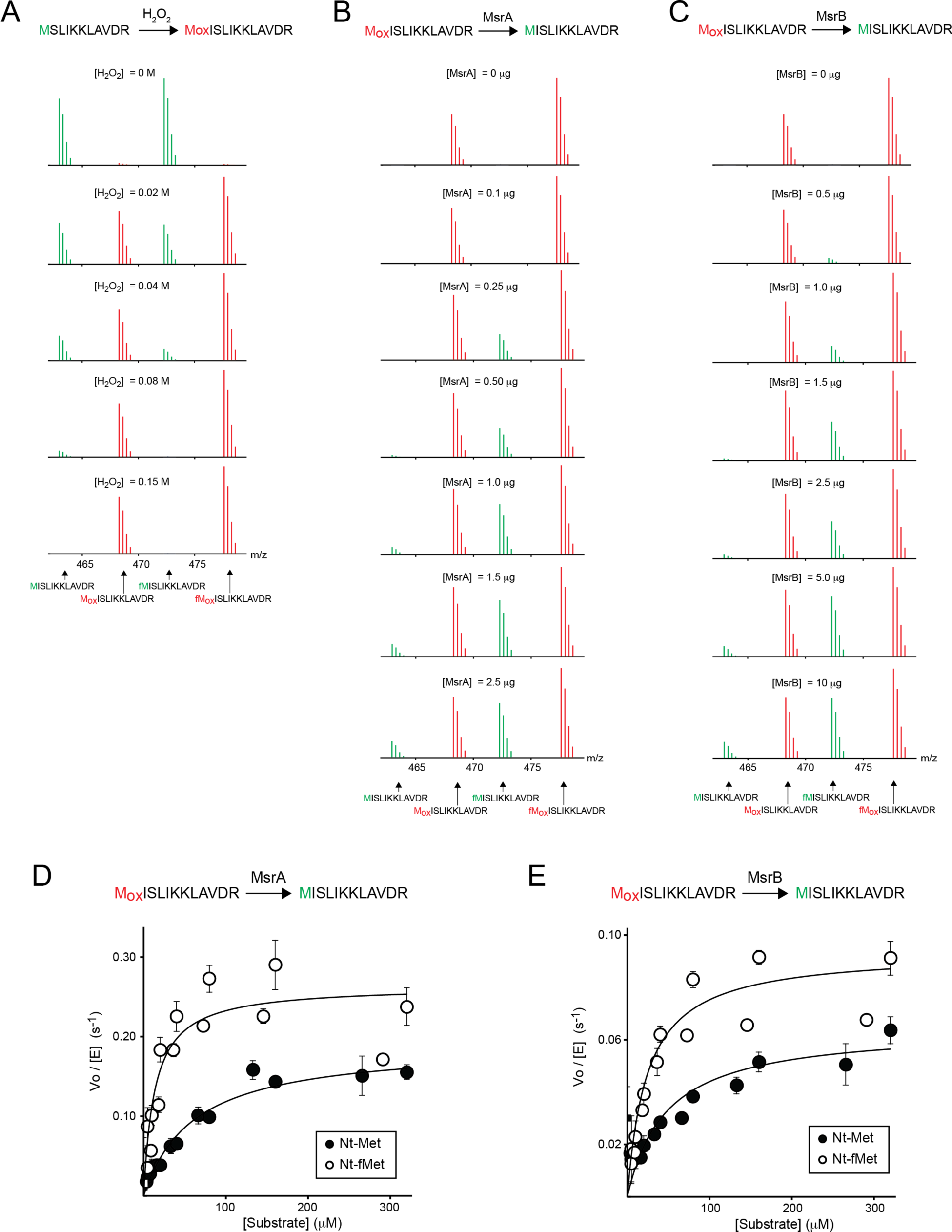
The effect of formylation on oxidation and reduction of Nt-Mets within the synthetic peptide MISLIKKLAVDR (Related to Figure 1, Table 1.) **A)** Nt-Met oxidation as a function of H_2_O_2_ concentration. **B)** Nt-Met reduction by MsrA as a function of enzyme concentration. **C)** Nt-Met reduction by MsrB as a function of enzyme concentration. The spectra illustrate the +3 charge state of the peptide. Green and red spectra distinguish the unoxidized and oxidized forms of the peptide, respectively. The expected m/z of unformylated/unoxidized, unformylated/oxidized, formylated/unoxidized and formylated/oxidized forms of the peptide are indicated below the spectra. **D)** Michaelis-Menten plots indicating the effect of formylation on the reduction kinetics of MsrA. **E)** Michaelis-Menten plots indicating the effect of formylation on the reduction kinetics of MsrB.

**Figure S2.**
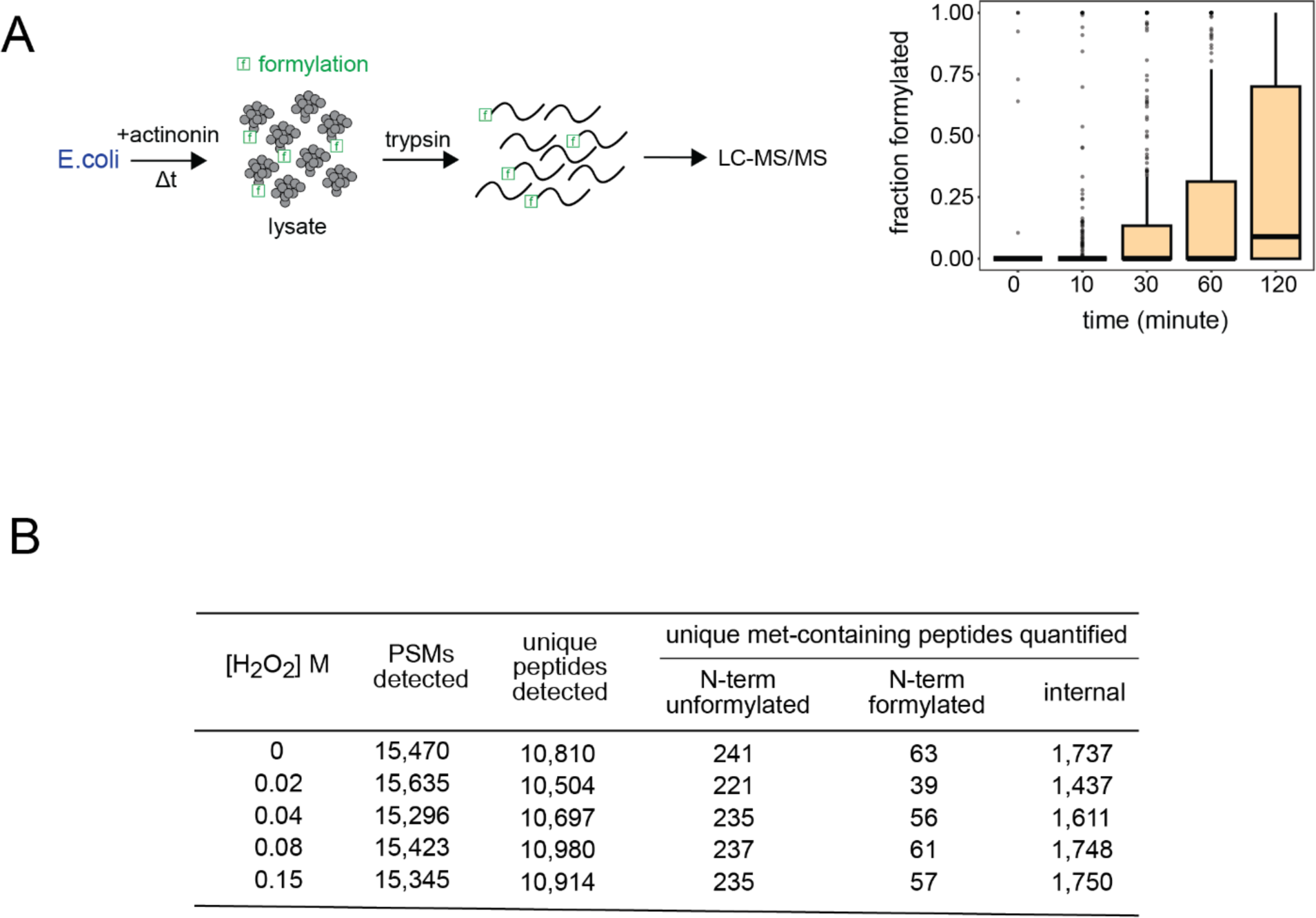
Accumulation of Nt-fMets in actinonin-treated *E. coli* and coverage of proteome-wide Nt-Met oxidation experiments (Related to Figure 2.) **A)** To obtain a population of Nt-fMet-containing peptides, *E.coli* cultures were treated with actinonin for variable durations. Cell extracts were then treated with trypsin and analyzed by LC-MS/MS. For each detected peptide that contained a N-terminal methionine, the fraction of the total signal intensity (of all modified forms) that was attributed to the formylated form of the peptide was measured. The box plots indicate distributions of fractional formylation for Nt-Met-containing peptides. The black lines within boxes are the median measurements, the whiskers indicate the entire range excluding outliers, and the dots indicate the outliers (>2 SD.) **B)** Coverage of proteome-wide oxidation experiments indicating numbers of detected methionine-containing peptides.

**Figure S3.**
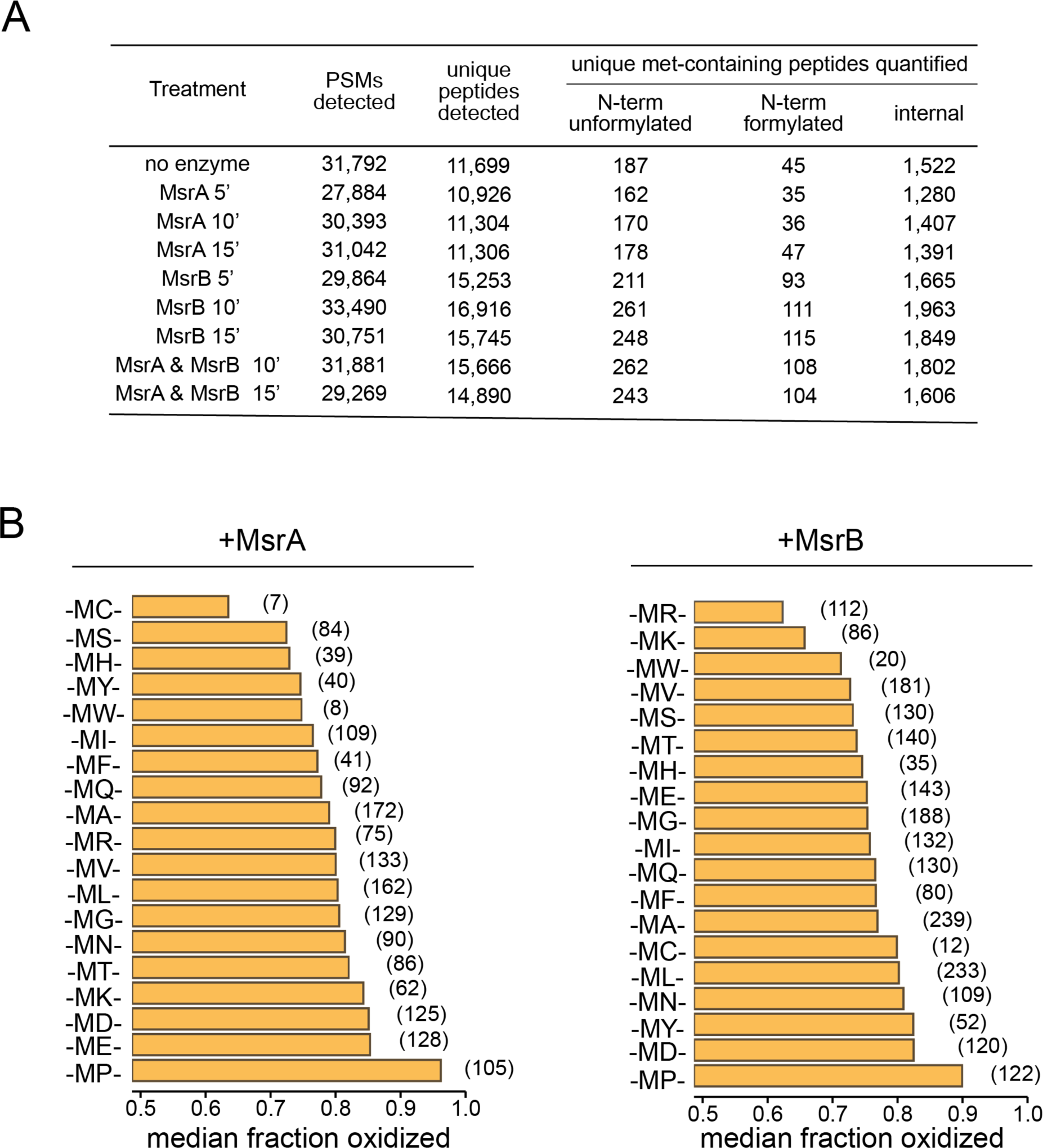
Coverage of proteome-wide reduction experiments on actinonin-treated extracts and C-side (M-AA_X_) neighboring residue effects on Nt-Met reduction efficiencies (Related to Figure 3.) **A)** Coverage of proteome-wide reduction experiments of actinonin-treated extracts, indicating numbers of detected methionine-containing peptides. **B)** Median fractional oxidation levels of subsets of peptides with the indicated neighboring residues on C-side (M-AA_X_) of Mets after exposure to MsrA or MsrB for 10 minutes.

**Figure S4.**
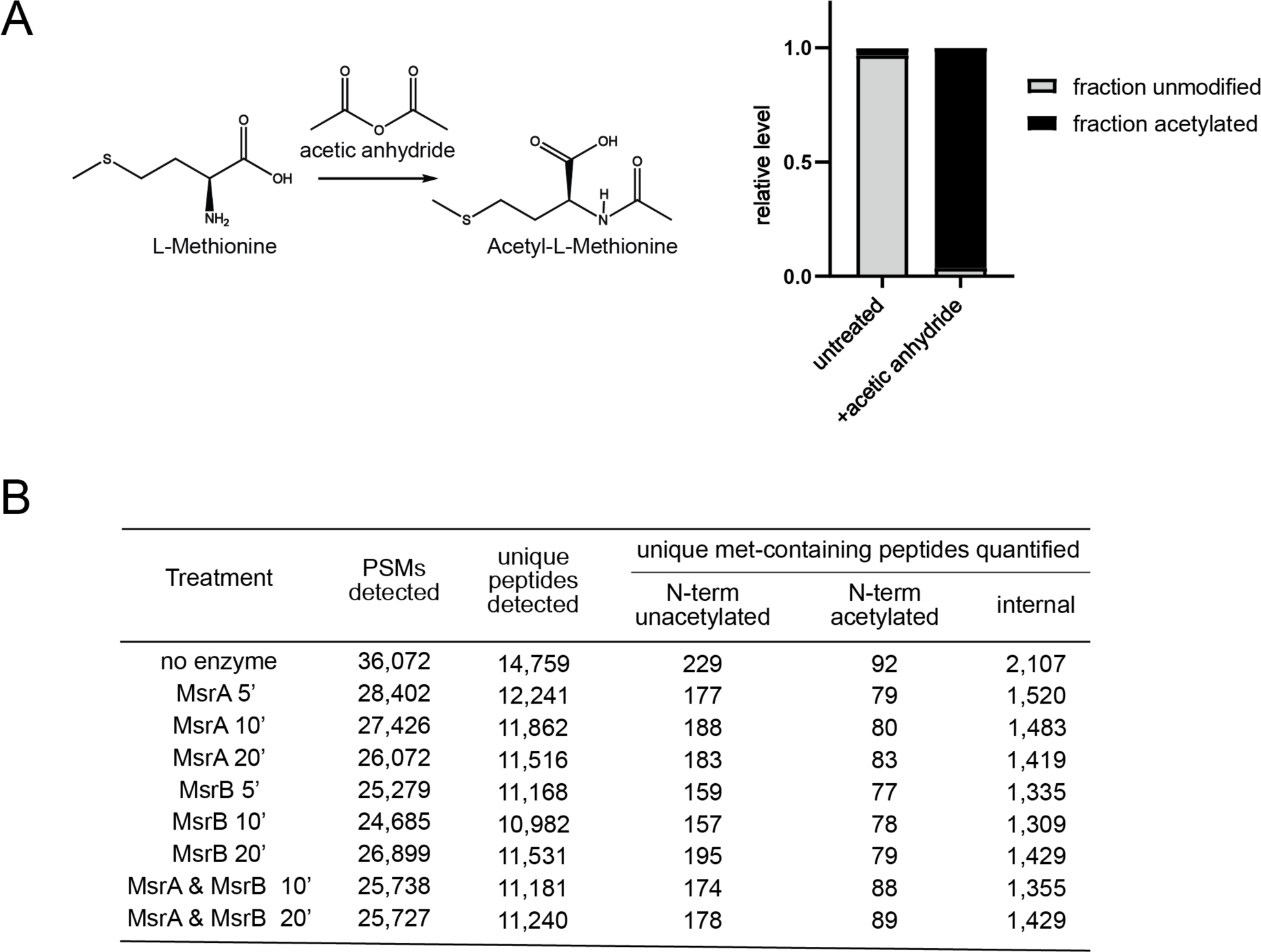
Efficiency of chemical acetylation of trypsinized *E. coli* extracts and coverage of proteome-wide reduction experiments on acetylated extracts (Related to Figure 4.) **A)** Based on the analysis of proteomic data, fraction of the total intenity of PSMs that were modified by N-acetylation with and without treatment with acetic anhydride. **B)** Coverage of proteome-wide reduction experiments of acetylated extracts, indicating numbers of detected methionine-containing peptides.

**Figure S5.**
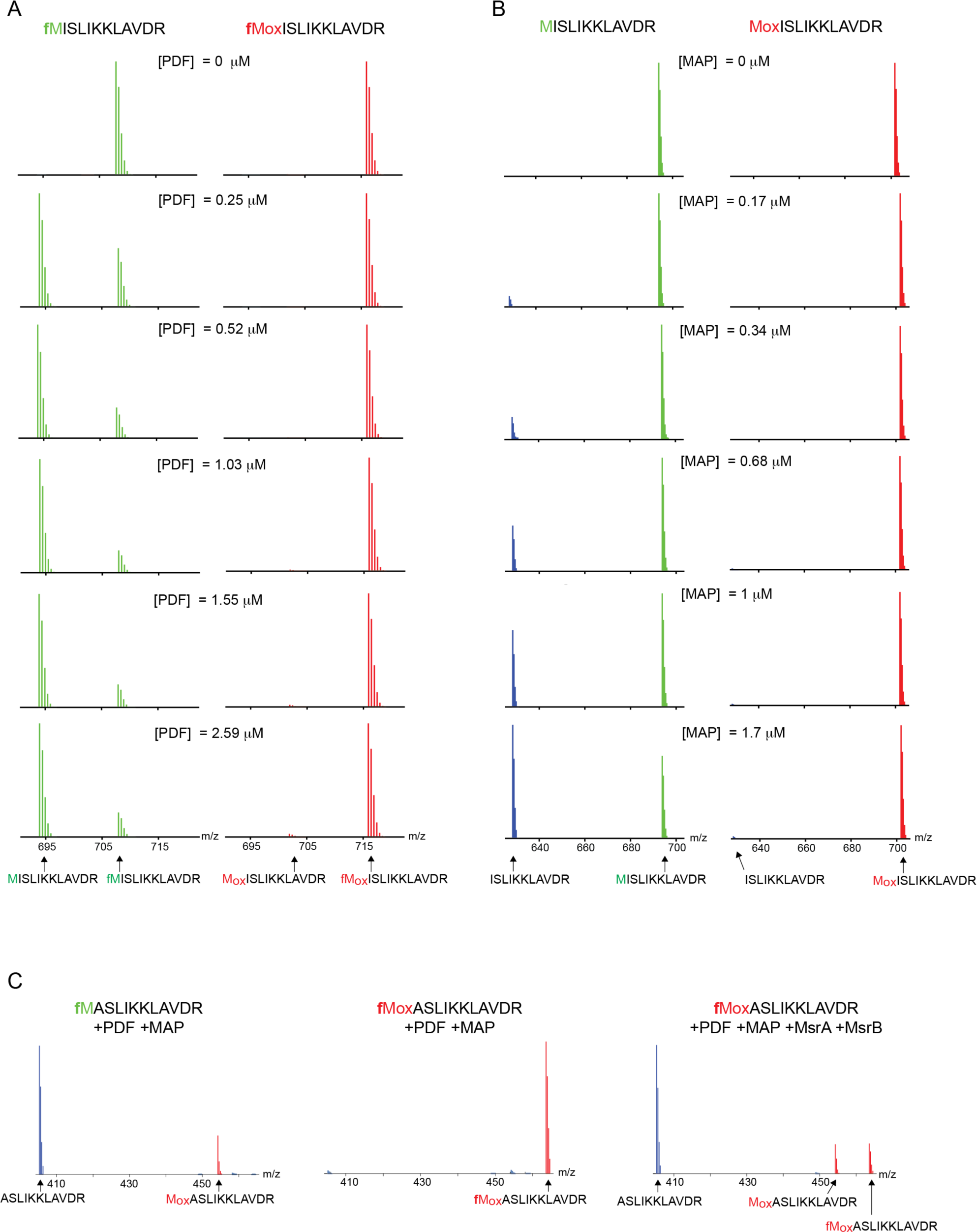
Mass spectra indicating the effect of oxidation on the deformylation and cleavage of Nt-Mets within the synthetic peptide MISLIKKLAVDR (Related to Figure 5.) **A)** Nt-fMet deformylation as a function of PDF concentration. The left and right columns of spectra were generated using unoxidized and oxidized Nt-fMet-containing peptides as substrates, respectively. **B)** Nt-Met cleavage as a function of MAP concentration. The left and right columns of spectra were generated using unoxidized and oxidized Nt-Met-containing peptides as substrates, respectively. **C)** Nt-fMet deformylation and subsequent cleavage by combined addition of PDF and MAP in the presence or absence of MsrA and MsrB. Unoxidized or oxidized peptides were used as substrates as indicated.

## Supplementary Tables

**Supplementary Table S1.** Proteome-wide analysis of oxidation efficiencies Met-containing peptides. (TableS1.xlsx)

**Supplementary Table S2.** Reduction efficiencies of Met-containing peptides in actinonin-treated proteomes by MsrA and MsrB. (TableS2.xlsx)

**Supplementary Table S3.** Reduction efficiencies of Met-containing peptides in acetylated proteomes by MsrA and MsrB. (TableS3.xlsx)

**Supplementary Table S4.** Comparison of PSM counts of unoxidized and oxidized Met-containing peptides oxidation levels between WT and *λ¢<fmt E. coli*.

